# Haploidy-linked cell proliferation defects limit larval growth in Zebrafish

**DOI:** 10.1101/2022.05.12.491746

**Authors:** Kan Yaguchi, Daiki Saito, Triveni Menon, Akira Matsura, Miyu Hosono, Takeomi Mizutani, Tomoya Kotani, Sreelaja Nair, Ryota Uehara

**Affiliations:** Graduate School of Life Science, Hokkaido University, Kita 21, Nishi 11, Kita-Ku, Sapporo, 001-0021, Japan; Faculty of Advanced Life Science, Hokkaido University, Kita 21, Nishi 11, Kita-Ku, Sapporo, 001-0021, Japan; Janelia Research Campus, Howard Hughes Medical Institute, Ashburn, Virginia, USA; Department of Life Science and Technology, Faculty of Engineering, Hokkai-Gakuen University, Minami 26, Nishi 11, Chuo-ku, Sapporo, 064-0926, Japan; Department of Biological Sciences, Faculty of Science, Hokkaido University, Kita 10, Nishi 8, Kita-Ku, Sapporo 060-0810, Japan; Department of Biosciences and Bioengineering, Indian Institute of Technology Bombay, Powai, Mumbai, 400076, India

**Author notes:** Corresponding authors: Sreelaja Nair, Department of Biosciences and Bioengineering, Indian Institute of Technology Bombay, Powai, Mumbai, 400076, India; Ryota Uehara, Faculty of Advanced Life Science, Hokkaido University, Kita 21, Nishi 11, Kita-Ku, Sapporo 001-0021, Japan. Kan Yaguchi and Daiki Saito are the co-first authors of this paper.

**Keywords:** Ploidy, Centrosome, Zebrafish

## Abstract

Haploid larvae in non-mammalian vertebrates are lethal with characteristic organ growth retardation collectively called “haploid syndrome.” In contrast to mammals whose haploid intolerance is attributed to imprinting misregulation, the cellular principle of haploidy-linked defects in non-mammalian vertebrates remains unknown. Here, we investigated cellular defects that disrupt the ontogeny of gynogenetic haploid zebrafish larvae. Unlike diploid control, haploid larvae manifested unscheduled cell death at the organogenesis stage, attributed to haploidy-linked p53 upregulation. Moreover, we found that haploid larvae specifically suffered the gradual aggravation of mitotic spindle monopolarization during 1-3 days post fertilization, causing spindle assembly checkpoint-mediated mitotic arrest throughout the entire body. High-resolution imaging revealed that this mitotic defect accompanied the haploidy-linked centrosome loss occurring concomitantly with the gradual decrease in larval cell size. Either resolution of mitotic arrest or depletion of p53 partially improved organ growth in haploid larvae. Based on these results, we propose that haploidy-linked mitotic defects and cell death are parts of critical cellular causes shared among vertebrates that limit the larval growth in the haploid state, contributing to an evolutionary constraint on allowable ploidy status in the vertebrate life cycle.

## Introduction

Compared to plants or fungi, the life cycle in animals is restricted regarding ploidy. In plant and fungal life cycles, cells undergo mitotic divisions and develop multicellular bodies in the haploid and diploid generations (Mable and Otto, 1998). In contrast, the multicellular stage in animals is restricted to the diploid generation, except in some orders of haplodiplontic invertebrates (Mable and Otto, 1998; Otto and Jarne, 2001). The ploidy restriction is especially stringent in vertebrates: Haploid embryos generated through egg activation without the contribution of one parent’s genome (e.g., parthenogenesis, gynogenesis, or androgenesis) are almost invariably lethal in vertebrates (Wutz, 2014).

In mammals, haploid embryonic lethality is mainly attributed to misregulation of imprinted genes. Differential expression of parental alleles of imprinted genes disables the development of mammalian uniparental haploid embryos beyond the preimplantation blastocyst stage (Leeb and Wutz, 2013; Tilghman, 1999). On the contrary, non-mammalian vertebrates are devoid of parent-specific genomic imprinting and, hence, free from imprinting-associated developmental defects. However, despite the lack of the detrimental feature of imprinting misregulation, haploid larvae are lethal even in non-mammalian vertebrates.

Compared to mammals, ontogenetic defects become evident at later developmental stages in haploid non-mammalian vertebrates. For example, haploid larvae in zebrafish undergo gastrulation and somite formation and initiate organogenesis by 24 hours post fertilization (hpf) with only a modest delay from diploid counterparts (Menon et al., 2020). Haploid larvae in other fish and amphibians also reach the organogenesis stage (Fankhauser, 1938; Hamilton, 1966; Uwa, 1965). However, after the onset of organogenesis, haploid larvae in these non-mammalian species manifest pleiotropic developmental defects and succumb to lethality after hatching. For example, haploid embryos and larvae suffer abnormalities in collective cell migration during gastrulation (Menon et al., 2020), water inflow control through ectoderm in gastrulae (Hamilton and Tuft, 1972), prechordal plate function (Araki et al., 2001), epidermal cell arrangements during post-gastrula stages (Ellinger, 1979; Ellinger and Murphy, 1980), melanophore pigmentation (Uwa, 1965), and blood circulation (Uwa, 1965). These developmental abnormalities have been collectively called “haploid syndrome” (Dasgupta and Matsumoto, 1972; Hamilton, 1966; Luo and Li, 2003).

In the complex profile of haploid syndrome, a particularly common abnormality across non-mammalian vertebrate species is severe growth retardation of organs, such as the brain and eyes (Dasgupta and Matsumoto, 1972; Fankhauser and Griffiths, 1939; Hertwig, 1911; Oppermann, 1913; Purdom, 1969; Uwa, 1965). Forced diploidization of haploid larvae by artificial induction of whole-genome duplication in the early cleavage stages resolves organ retardations, suggesting that the haploid state per se, rather than loss of heterozygosity of deleterious recessive alleles, causes these defects (Menon and Nair, 2018; Nagy et al., 1978; Streisinger et al., 1981; Subtelny, 1958). Though previous studies identify global developmental errors associated with haploidy, the specific cell biological triggers of haploid organ retardations remain largely unknown. Elucidation of a fundamental cellular cause of the haploidy-linked organ retardations in non-mammalian vertebrates would provide a profound insight into evolutionary constraints that limit the multicellular stage of vertebrate life cycles exclusively to the diploid generation.

In this study, we investigated cellular defects in haploid zebrafish during the larval stages, when morphological abnormalities became evident (Kroeger et al., 2014). Haploid larvae suffered unscheduled cell death associated with p53 upregulation. High-resolution imaging revealed that, as cell size gradually decreased after 12 hpf, drastic centrosome loss occurred almost exclusively in haploid cells below a certain size threshold. This cell size-linked centrosome loss led to the gradual accumulation of mitotic defects in haploid larvae, which coincided with the characteristic temporal pattern of organ growth retardation at the developmental stage. Alleviating mitotic arrest or artificially reducing p53 levels partially improved organ growth, indicating that haploidy-linked poor organ growth stems at least in part from these cellular defects. Our results revealed haploidy-linked incoordination of cell proliferation, whose progression pattern may shape the characteristic profile of “haploid syndrome” in non-mammalian vertebrates.

## Results

### A drastic increase in unscheduled apoptosis in haploid larvae

We investigated the developmental processes of haploid and diploid zebrafish larvae generated by *in vitro* fertilization using UV-irradiated and non-irradiated spermatozoa, respectively (Fig. S1). At 3.5 days post fertilization (dpf), by when most organs have formed, haploid larvae manifested typical “haploid syndrome” defects, such as curled, short body axis and reduced brain and eye sizes compared with diploid counterparts (Fig. 1A and B) (Kroeger et al., 2014; Luo and Li, 2003; Purdom, 1969; Uwa, 1965). The morphological defects of haploid syndrome occurred in varying phenotypic grades, with some larvae manifesting severe defects and others with milder defects within a clutch (Fig. 1A). Consistent with this, we observed a larger variance of body and organ size distributions in haploid larval groups than in diploids (Fig. 1B). DNA content analysis using flow cytometry showed that the 1C and 2C populations (possessing one and two genome copies, respectively) predominated in the haploid larval groups at 3 and 5 dpf, confirming that cells in haploid zebrafish larvae retained the haploid state throughout the time duration (Fig. S1). This is in sharp contrast to haploid mammalian embryonic cells that quickly convert to diploids during early embryogenesis (Leeb and Wutz, 2011; Sagi et al., 2016).

**Figure 1.**
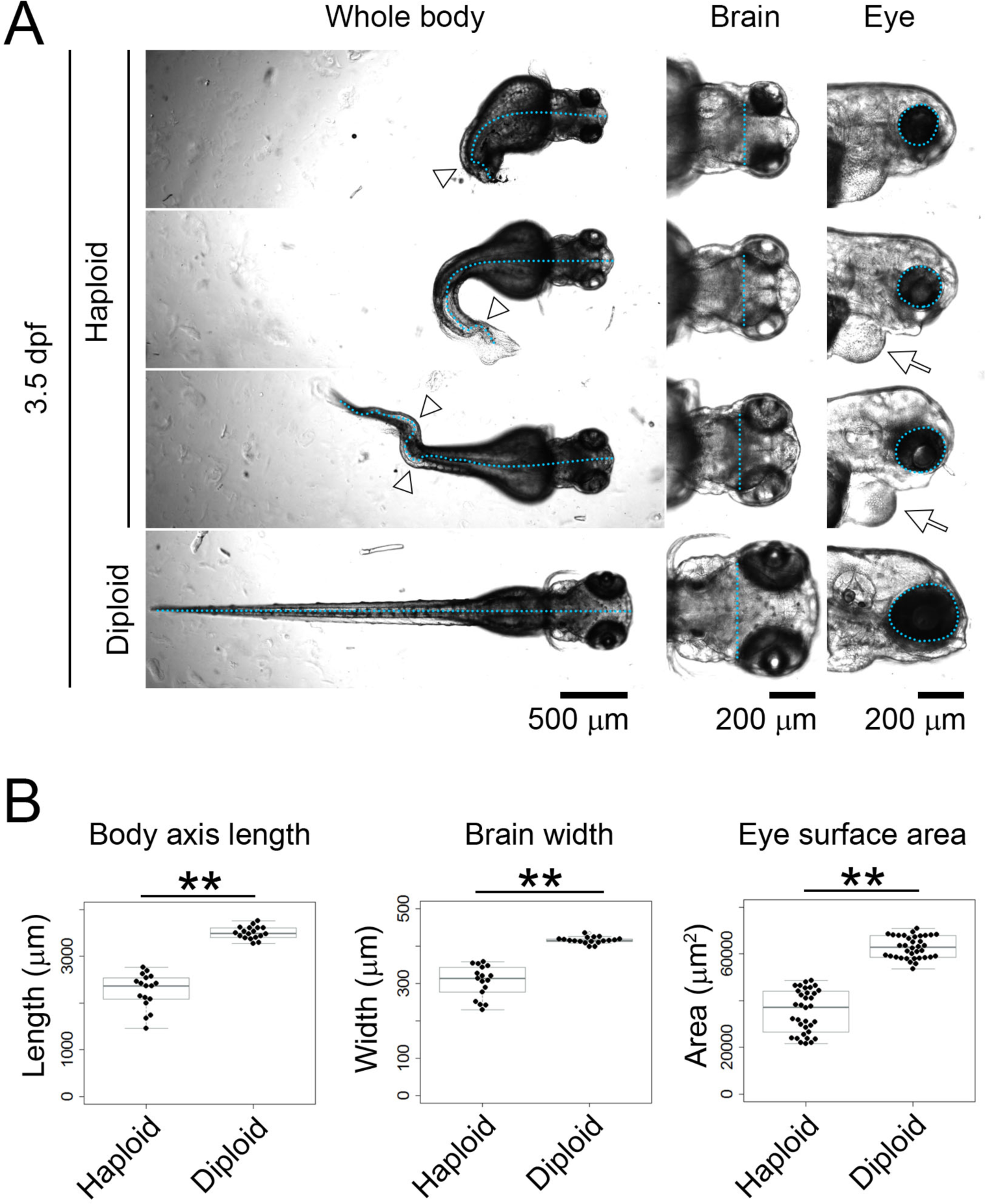
Morphological comparison between haploid and diploid larvae. **(A)** Transparent microscopy of haploid or diploid larvae at 3.5 dpf. Broken line: body axis (left panels), brain width (middle panels), or lateral eye contour (right panels). Arrowheads: curling or bending of body axis. Arrows: Edema. **(B)** Quantification of body axis length, brain width, or lateral eye area of haploid or diploid larvae in A. Box plots and beeswarm plots of ≥17 larvae (≥34 eyes) from three independent experiments (\*\**p* < 0.01, two-tailed *t*-test).

The drastically poor organ growth in haploid larvae indicated a possibility that haploid cells suffered defects in cell proliferation. To test this idea, we visualized apoptotic and mitotic cells by immunostaining cleaved caspase-3 and phospho-histone H3 (pH3), respectively (Mendieta-Serrano et al., 2013; Sorrells et al., 2013), in 3 dpf haploid and diploid larvae (Fig. 2A). In diploid larvae, cleaved caspase-3-positive cells were infrequent and found at specific sites, such as the optic tectum, mid-brain, and the inner retinal layers (Fig. 2A, B, and S2), reflecting programmed apoptosis underlying tissue organization (Yamashita, 2003). On the other hand, in haploid larvae, cleaved caspase-3-positive cells were frequently detected throughout the whole body (Fig. 2A, B, and S2). Quantification of the density of cleaved caspase-3-positive cells within the right midbrain demonstrated a significantly higher frequency of apoptotic cells in haploids than in diploids (Fig. 2C). Moreover, haploid larvae showed a significant increase in frequency of pH3-positive mitotic cells compared to diploids (Fig. 2A-C, see also Fig. S4E for 3-d distribution of pH3-positive cells in haploids). These results demonstrate that haploid larvae suffer irregularly patterned apoptosis and cell division throughout the body, potentially impairing organ growth during development. Therefore, we decided to address the molecular and cellular bases of these haploidy-linked defects in cell survival and proliferation.

**Figure 2.**
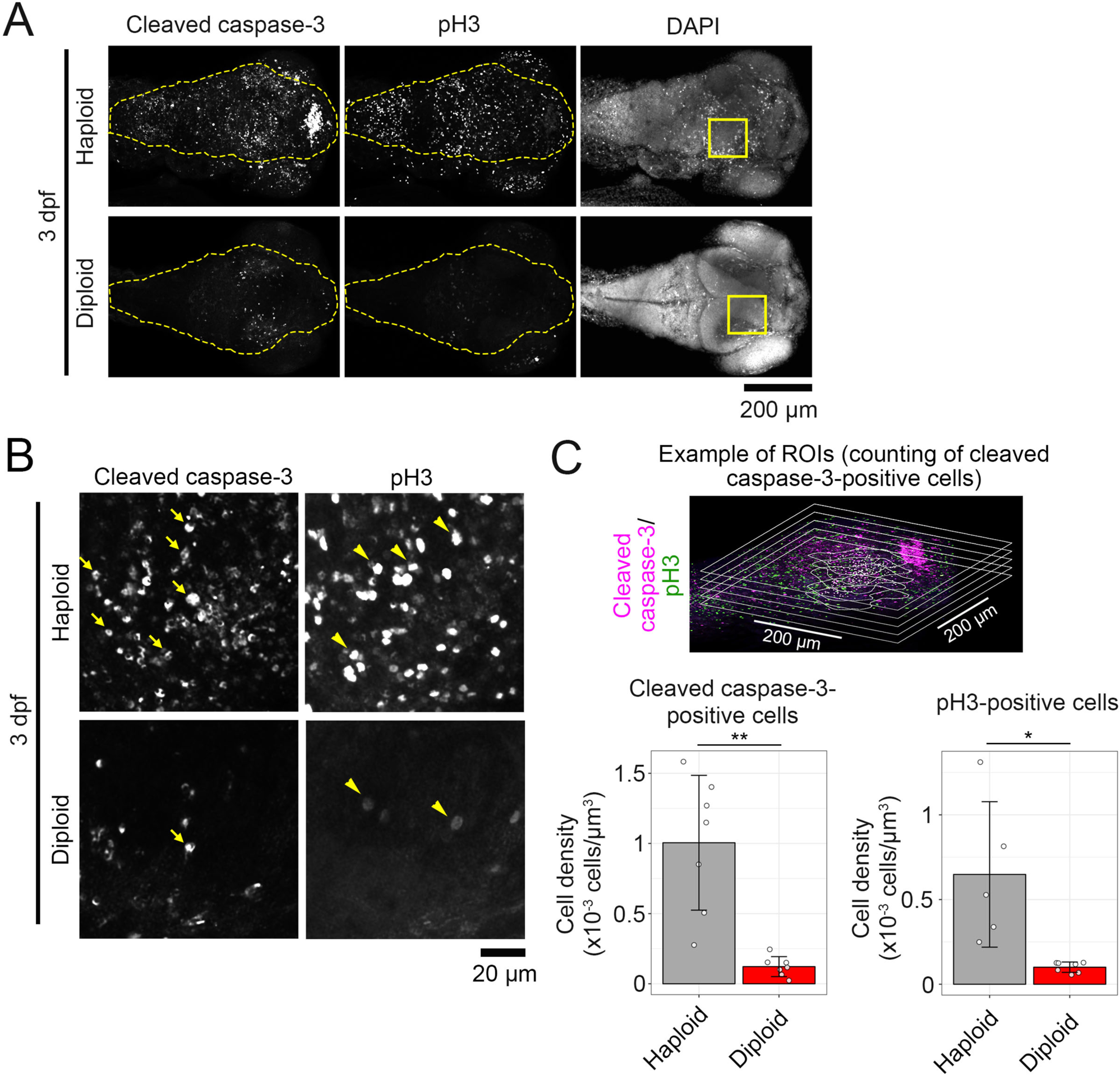
Haploid zebrafish larvae suffer irregularly increased apoptosis. **(A)** Immunostaining of cleaved caspase-3 and phospho-histone H3 (pH3) in whole-mount haploid or diploid larvae at 3 dpf. DNA was stained by DAPI. Z-projected images of confocal sections containing peripheral brain surface are shown. Broken lines indicate brain area. Boxes indicate the areas enlarged in *B*. **(B)** Enlarged views of haploid or diploid larvae in A. Arrows or arrowheads indicate examples of cleaved caspase-3- or pH3-positive cells, respectively. (**C**) The density (cell number per volume) of cleaved caspase-3- or pH3-positive cells in the right midbrain in haploid or diploid larvae in B. Examples of regions of interest and counted cell positions are shown on the top. Five or 8 confocal slice sections from the top of the brain were used for counting the number of caspase-3- or pH3-positive cells, respectively. Means ± standard deviation (SD) of ≥5 larvae from 2 independent experiments for haploids and diploids (\**p* < 0.05, \*\**p* < 0.01, two-tailed *t*-test). All larvae observed in this experiment are shown in Fig. S2.

### Irregular p53 upregulation limits organ growth in haploid larvae

We next sought to determine the cause of the haploidy-linked unscheduled apoptosis. As a candidate apoptosis inducer, we investigated expression levels of p53 protein in haploid and diploid larvae at 3 dpf by immunoblotting (Fig. 3A). p53 expression was significantly higher in haploid larvae than in diploids (Fig. 3A and B). p53 level in haploids was even higher than that in diploid larvae irradiated with UV to induce DNA damage-associated p53 upregulation (Fig. 3B).

**Figure 3.**
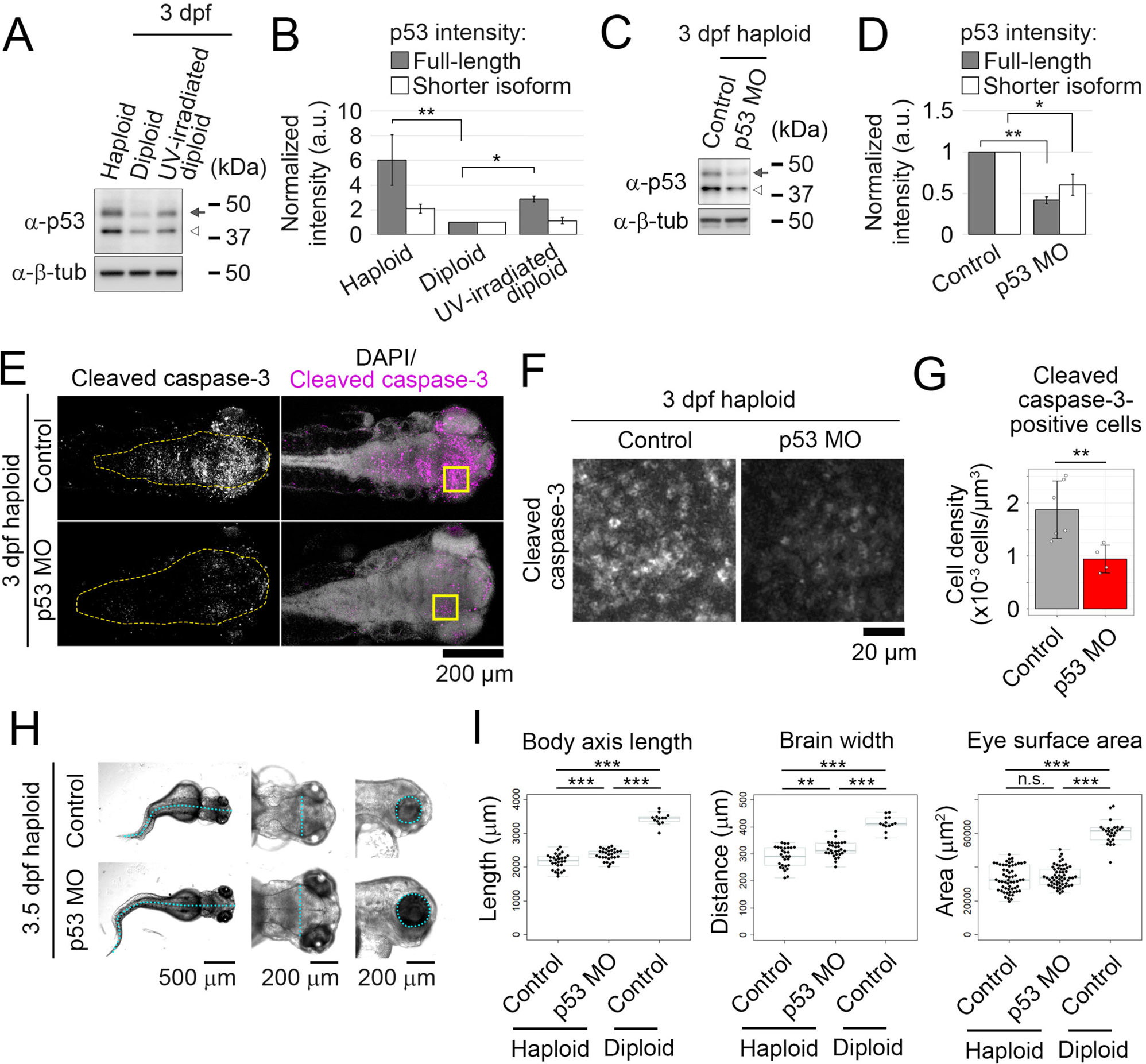
p53 upregulation limits organ growth in haploid larvae. **(A, C)** Immunoblotting of p53 in haploid, diploid, or UV-irradiated diploid larvae (A) or haploid larvae treated with control or p53 antisense morpholino (C) at 3 dpf. Arrows and open arrowheads indicate full-length and shorter p53 isoforms, respectively. β-tubulin was detected as a loading control. **(B**, **D)** Quantification of relative expression of p53 proteins in A (B) or C (D). Mean ± standard error (SE) of 3 independent experiments (\**p* < 0.05, \*\**p* < 0.01, the ANOVA followed by the post-hoc Tukey test in B, and two-tailed t-test in D). **(E)** Immunostaining of active caspase-3 in whole-mount haploid control or p53 morphant at 3 dpf. DNA was stained by DAPI. Z-projected images of confocal sections containing peripheral brain surfaces are shown. Broken lines indicate brain area. Boxes indicate the areas enlarged in *F*. **(F)** Enlarged views of morphants in E. **(G)** The density of cleaved caspase-3-positive cells in the right midbrain in morpholino-injected haploid larvae in F. Two confocal slice sections from the top of the brain were used for cell counting. Means ± SD of 6 control morphants and 4 p53 morphants from 2 independent experiments (\*\**p* < 0.01, two-tailed *t*-test). All larvae observed in this experiment are shown in Fig. S3A. **(H)** Transparent microscopy of haploid control and p53 morphants at 3.5 dpf. Broken line: body axis (left panels), brain width (middle panels), and lateral eye contour (right panels). **(I)** Quantification of the body axis, brain width, and lateral eye area in haploid control or p53, or diploid control morphants at 3.5 dpf. Box plots and beeswarm plots of ≥13 larvae (≥26 eyes) from 3 independent experiments. Asterisks indicate statistically significant differences among samples (n.s.: not significant, \*\**p* < 0.01, \*\*\**p* < 0.001, the ANOVA followed by the post-hoc Tukey test).

To investigate the causality between irregular p53 upregulation and haploidy-linked defects, we next tested the effects of morpholino-mediated p53 depletion on apoptosis in haploid larvae. p53 expression was substantially reduced by a p53-specific morpholino (Fig. 3C and D). In control haploid morphants injected with a 4-base mismatch morpholino, cleaved caspase-3-positive apoptotic cells were detected throughout the body (Fig. 3E and S3A). Depletion of p53 significantly reduced the frequency of cleaved caspase-3-positive cells in haploid larvae compared to control (Fig. 3F and G). DNA content in haploid p53 morphants was equivalent to that in control haploids, demonstrating that the ploidy level did not change upon p53 depletion in haploid larvae (Fig. S3B). These results suggest that p53 upregulation contributes to enhanced apoptosis in haploid larvae.

We also tested the effect of p53 depletion on organ size in haploid larvae (Fig. 3H). p53 depletion significantly increased body-axis length and brain width assessed at the mid-brain in haploid larvae (Fig. 3H and I), while the size increase for body axis length, brain width, or eye surface area was about 15.9%, 22.3%, or 5.0%, respectively, of the difference between haploids and diploids. p53 depletion notably eliminated the population of haploid larvae that suffered particularly severe organ size reduction, suggesting modest alleviation of the haploidy-linked poor organ growth (Fig. 3I). The above data indicate p53-induced apoptosis as a part of the cause of haploidy-linked organ growth retardation.

### Haploidy larvae suffer severe mitotic arrest

Next, we sought to understand the reason for the irregular cell proliferation in haploid larvae and its possible influence on developmental defects in the haploid state. To quantitively analyze the haploidy-linked change in cell proliferation pattern, we conducted flow cytometry for DNA content and pH3-positive cell proportion in 1-3 dpf haploid and diploid whole-larval cells (Fig. 4A and B). In diploid larvae, the proportion of pH3-positive cells was 1.6 % at 1 dpf, which decreased to 0.4% at 3 dpf (Fig. 4A and B), possibly reflecting the transition from proliferative to postmitotic state in different cell lineages during larval stages (Li et al., 2000; Sugiyama et al., 2009). On the other hand, the proportion of pH3-positive cells in haploid larvae was 1.6% at 1 dpf and 1.2% at 3 dpf (Fig. 4A and B). The mitotic index was significantly higher in haploid larvae than in diploids at 2-3 dpf (Fig. 4B), consistent with the finding in the whole-mount larval imaging (Fig. 2A-C).

**Figure 4.**
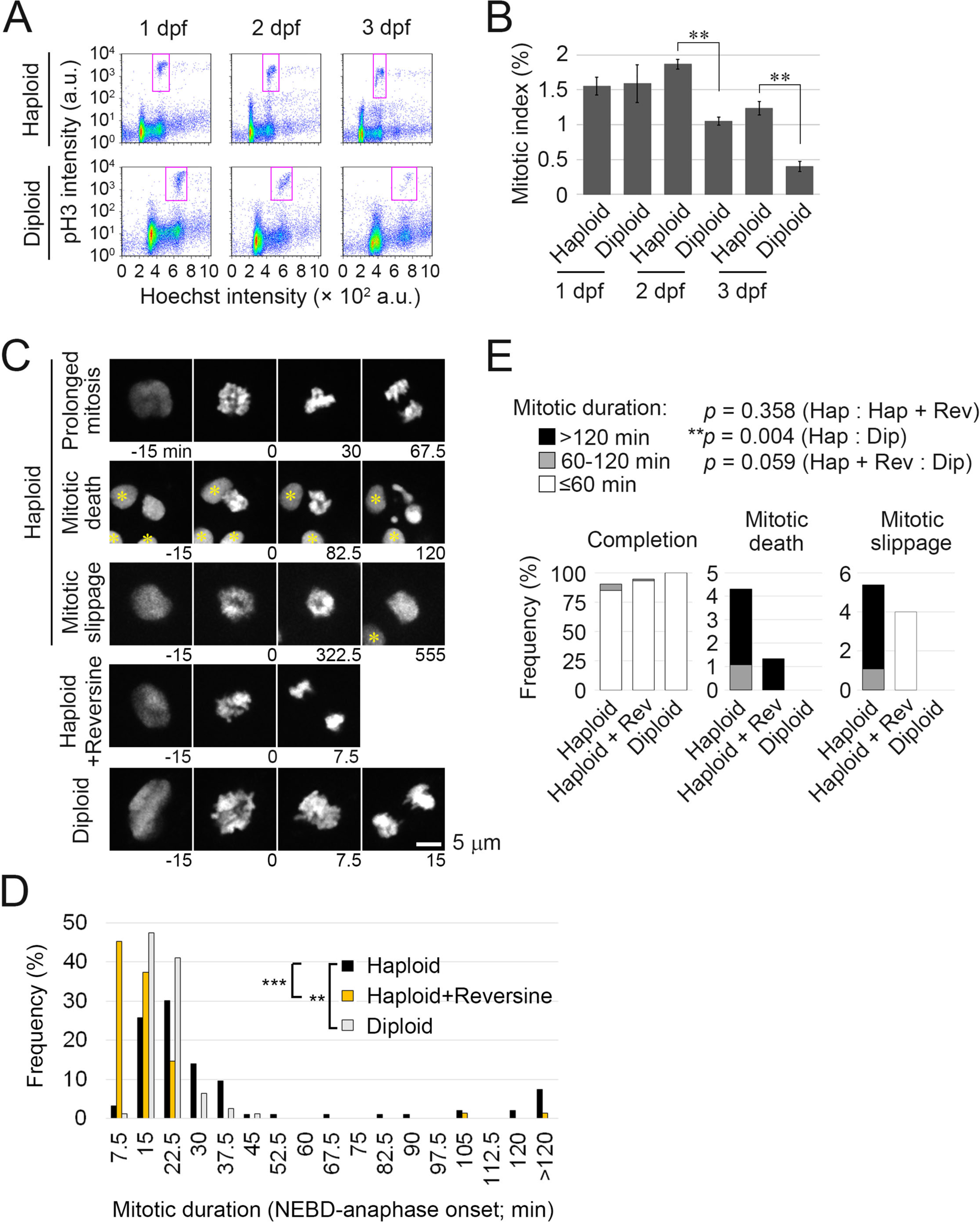
Frequent mitotic delay and failures in haploid larvae. **(A)** Flow cytometric analysis of DNA content (Hoechst signal) and mitotic proportion (marked by anti-pH3) in isolated haploid or diploid larval cells at 1, 2, or 3 dpf. Magenta boxes indicate the pH3-positive mitotic populations. **(B)** Quantification of mitotic index in A. Mean ± SE of ≥3 independent experiments (\*\**p* < 0.01, two-tailed *t*-test). **(C)** Live images of endothelial cells expressing histone H2B-mCherry in haploid, diploid, or reversine-treated haploid larvae. Images were taken at a 7.5 min interval from 1.5 to 3 dpf. Reversine was administrated approximately from 1.5 dpf. Asterisks indicate neighbor cells. **(D)** Distribution of mitotic length (time duration from NEBD to anaphase onset) in C. At least 75 cells of 6 larvae from 6 independent experiments were analyzed. Asterisks indicate statistically significant differences among samples (\*\**p* < 0.01, \*\*\**p* < 0.001, the ANOVA followed by the post-hoc Tukey test; among cells in the “>120 min” bin in the histogram, the ones whose mitotic exit time could not specified were excluded from the statistic analysis). **(E)** Frequency of mitotic fates in C. Data were sorted into separated graphs by mitotic fates (completion, mitotic death, or mitotic slippage). At least 75 cells of 6 larvae from 6 independent experiments were analyzed. *P*-values obtained by the Fisher exact test with the Benjamini-Hochberg multiple testing correction are shown at the top.

To address the cause of the mitotic index increase in haploids, we conducted live imaging and analyzed the mitotic progression of endothelial cells in 1.5-3 dpf haploid and diploid larvae stably expressing histone-H2B-mCherry (Tg(fli1:h2b-mCherry)/ncv31Tg) (Yokota et al., 2015). In diploid larvae, almost all cells that entered mitosis (marked by nuclear envelope breakdown (NEBD)) completed chromosome alignment and segregation within 30 min after mitotic entry (Fig. 4C and D). On the other hand, 15% of haploid cells spent > 60 minutes in the mitotic phase, revealing a severe mitotic delay in haploid larvae. Such mitotically delayed haploid cells often underwent mitotic death or mitotic slippage (exiting mitosis without proper chromosome segregation) (Fig. 4C and E). Mitotic progression was slower in haploid larvae than in diploids for the duration of assessment from 1.5-3 dpf, with more frequent mitotic defects later in the observation (Fig. S4A-D). We also observed severe chromosome misalignment in haploid cells delayed in mitosis (Fig. 4C), suggesting that the increased mitotic index and mitotic delay seen in haploid larvae could be due to the activation of the spindle assembly checkpoint (SAC).

### Gradual aggravation of mitotic spindle monopolarization with centriole loss in haploid larvae during 1-3 dpf

To understand the cause of the frequent mitotic arrest with chromosome misalignment in haploid larvae, we investigated mitotic spindle organization by immunostaining of α-tubulin and centrin, which mark microtubules and the centrioles, respectively, in the head region of haploid and diploid larvae from 0.5 to 3 dpf (Fig. 5 and S5). Almost all mitotic cells in diploids possessed bipolar spindles with the normal number of 4 centrioles in all stages tested. At 0.5 dpf, haploid embryonic cells also possessed bipolar spindles with 4 centrioles (Fig. 5A-C, and S5A-D). However, as development progressed, cells in haploid larvae had monopolar spindles with reduced centriole number and severe chromosome misalignment (Fig. 5B and C). In the head region of haploid larvae, including eyes, brain, skin, and olfactory organ, 14.5% of mitotic cells had monopolar spindles at 1 dpf, which increased to 32.6% and 61.5% at 2 and 3 dpf, respectively (Fig. 5B). The timing of the gradual aggravation of spindle monopolarization and centrosome loss corresponded well with the time when mitotic index became significantly higher in haploids than in diploids with increased cell death (Fig. 4A, B and S4D). The trajectory of centriole loss and spindle monopolarization was organ-specific: these defects commenced earlier or were more drastic in the eyes and brain than in the skin (Fig. S5A-D). However, the frequency of monopolar mitotic cells with centriole loss reached over 50% in all these organs by 3 dpf, revealing the general nature of the cellular defects in haploid larvae.

**Figure 5.**
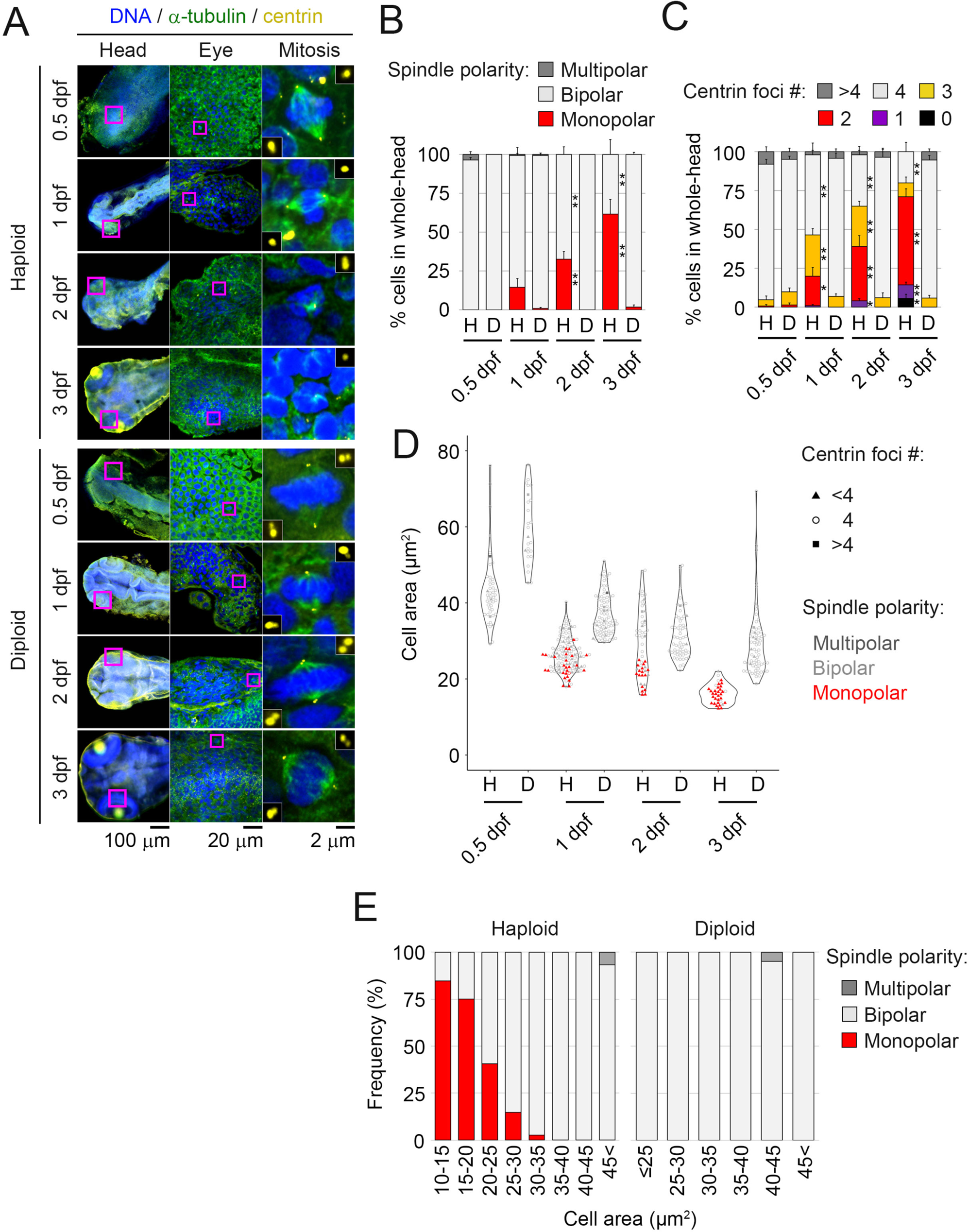
Centrosome loss and spindle monopolarization in haploid larvae. **(A)** Immunostaining of α-tubulin and centrin in haploid and diploid larvae at 0.5, 1, 2, and 3 dpf. Magenta boxes in the left or middle panels indicate the enlarged regions of eyes (shown in the middle panels) or mitotic cells (shown in the right panels), respectively. Insets in the right panels show 3× enlarged images of centrioles. **(B**, **C)** Frequency of spindle polarity and centrin foci number in mitotic cells in the whole-head region (including eyes, brain, skin epithelia, and olfactory organ) at different developmental stages. Mean ± SE of ≥4 larvae from ≥2 independent experiments (≥98 cells were analyzed for each condition; asterisks indicate statistically significant differences from diploids at the corresponding time points; \**p* < 0.05, \*\**p* < 0.01, two-tailed *t*-test). Data points taken from the eyes, the brain, or skin epithelia are also shown separately in Fig. S5. **(D**, **E)** Areas of the largest confocal section of subepidermal mitotic cells in haploid and diploid larvae during 0.5-3 dpf (D), or frequency of monopolar spindle in haploid and diploid larval cells sorted by their size (E). Centriole number and spindle polarity in each cell are indicated as depicted in the graph legends in D. At least 27 cells from ≥4 larvae from ≥2 independent experiments were analyzed for each condition in D. At least 13 cells were analyzed for each cell size bin in E.

Since the first division, embryonic cell size continuously reduces through successive cell divisions during early development (Menon et al., 2020). Cell size reduction continued at the developmental stage when we observed the gradual aggravation of the centrosome loss in haploid larvae. Therefore, we tested the relationship between cell size and centriole number in mitotic cells in subepidermal cell layers in the head region of 0.5-3 dpf haploid or diploid larvae (Fig. 5D and E; *see Material and methods*). During 0.5-3 dpf, the largest slice area of mitotic cells reduced from 43 μm^2^ to 16 μm^2^ or from 61 μm^2^ to 29 μm^2^ in haploids or diploids, respectively (Fig. 5D). Interestingly, monopolar spindles with a reduced number of the centrioles were observed almost exclusively in the cells whose largest slice area was less than 35 μm^2^, suggesting a lower limit of cell size for supporting centrosome number homeostasis in haploid larvae (Fig. 5D and E). Below the limit, frequency of monopolar spindles monotonically increased as cell size reduced in haploids (Fig. 5E). In contrast, diploid larval cells whose size became lower than this limit still formed bipolar spindles. The above data indicates that the stage-specific aggravation of mitotic spindle monopolarization through the haploidy-specific centrosome loss results in severe mitotic arrest in haploid larvae.

### Resolution of SAC-dependent mitotic arrest improves larval growth in the haploid state

Finally, we sought to address the causality between haploidy-linked mitotic arrest and larvae growth defects. For this, we attempted to resolve the haploidy-linked mitotic arrest by suppressing SAC using reversine, an inhibitor of Mps1 kinase required for SAC activation (Santaguida et al., 2010). In live imaging of histone-H2B-mCherry, 97% of reversine-treated haploid larval cells underwent chromosome segregation or exited mitosis within 30 min after NEBD (Fig. 4C, D, and S4C). Notably, mitotic cell death was substantially suppressed by reversine (Fig. 4E). Therefore, the SAC inactivation resolved the severe mitotic delay and mitigated mitotic cell death in haploid larvae.

We next tested whether SAC inactivation resolved the abnormal cell proliferation pattern in haploid larvae. For this, we analyzed the mitotic index in 3 dpf haploid larvae treated with reversine using flow cytometry (Fig. 6A and B). Reversine significantly reduced the proportion of pH3-positive cells in haploid larvae to levels equivalent to that in diploids (Fig. 6A and B, see also Fig. 4A and B). In immunostained haploid larvae, we also observed a reduction in pH3-positive cells upon reversine treatment (Fig. 6C-E). These results indicate that the abnormal cell proliferation pattern with the high mitotic index in haploid larvae is mainly due to the SAC-dependent mitotic arrest.

**Figure 6.**
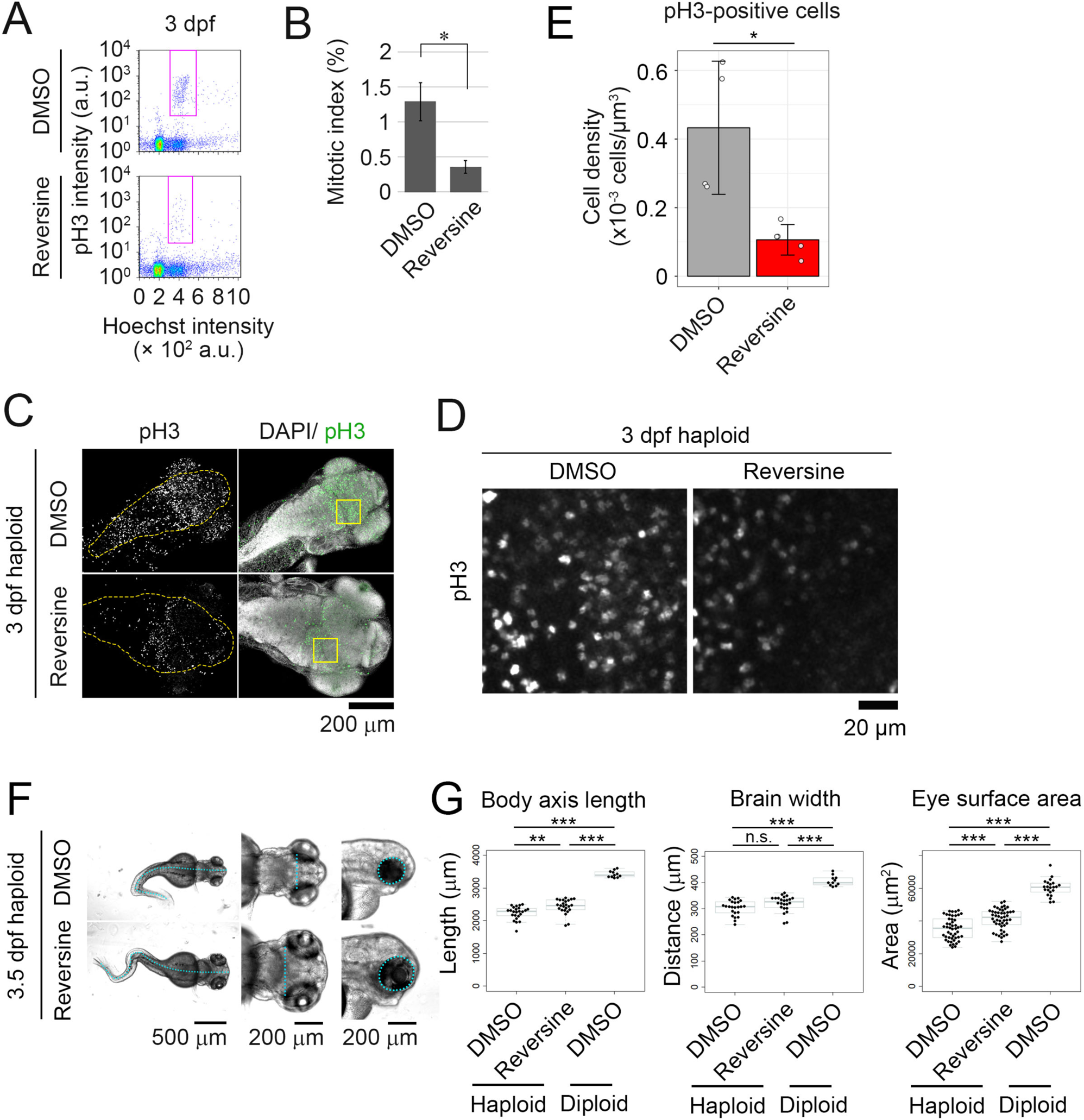
SAC inactivation mitigates abnormal mitotic patterns and organ growth defects in haploid larvae. **(A)** Flow cytometric analysis of DNA content (Hoechst signal) and mitotic proportion (marked by anti-pH3) in the cells isolated from 3-dpf haploid larvae treated with DMSO or reversine. DMSO or reversine was treated from 0.5 to 3 dpf. Magenta boxes indicate the pH3-positive mitotic populations. **(B)** Quantification of mitotic index in A. Mean ± SE of 4 independent experiments (\**p* < 0.05, two-tailed *t*-test). **(C)** Immunostaining of pH3 in haploid 3-dpf larvae, which were treated with DMSO or reversine from 1.5 to 3 dpf. Z-projected images of confocal sections containing peripheral brain surfaces are shown. Broken lines indicate brain area. Boxes indicate the areas enlarged in *D*. Note that pH3-positive cells in the immunostaining image are apparently more frequent than those estimated by flow cytometry, presumably because mitotic cells tended to localized at the surface region of larvae (see Fig. S4E for a vertical section of immunostained larvae). **(D)** Enlarged views of haploid or diploid larvae in *C*. (**E**) The density of pH3-positive cells in the right midbrain in haploid larvae in *C*. Eight confocal slice sections from the top of the brain were used for cell counting. Means ± SD of ≥4 larvae from 2 independent experiments (\**p* < 0.05, two-tailed *t*-test). **(F)** Transparent microscopy of haploid 3.5-dpf larvae treated with DMSO or reversine from 1.5 to 3.5 dpf. Broken line: body axis (left panels), brain width (middle panels), and lateral eye contour (right panels). **(G)** Measurement of the body axis, brain width, and lateral eye area in DMSO- or reversine-treated haploid, or DMSO-treated diploid 3.5-dpf larvae. Box plots and beeswarm plots of ≥11 larvae (≥22 eyes) from 3 independent experiments. Asterisks indicate statistically significant differences among samples (n.s.: not significant, \*\**p* < 0.01, \*\*\**p* < 0.001, the ANOVA followed by the post-hoc Tukey test).

Based on the finding that SAC inactivation mitigated abnormal cell proliferation in haploid larvae, we tested the effect of reversine treatment on organ growth in haploid larvae. Reversine treatment significantly increased body axis length and eye size in haploid larvae compared to DMSO-treated control (Fig. 6F and G), while the size increase for body axis length, brain width, or eye surface area was about 16.4%, 15.8%, or 23.8%, respectively, of the difference between haploids and diploids. These results indicate that the haploidy-linked mitotic stress with SAC activation is a part of the cause of organ growth defects in haploid larvae.

## Discussion

Many reports have revealed tissue-level developmental abnormalities in haploid embryos since the discovery of haploid embryonic lethality in vertebrates more than 100 years ago (Hertwig, 1911). However, there have been surprisingly few descriptions of cellular abnormalities underlying these tissue defects, precluding an evidence-based understanding of cellular principles that limit developmental ability in haploid embryos. Our basic descriptions of haploidy-linked cellular defects in haploid zebrafish larvae would provide a frame of reference for why vertebrates can not develop in the haploid states.

### Cell proliferation defects limit organ growth in haploid zebrafish larvae

Poor organ growth is a common feature of haploid syndrome in non-mammalian vertebrates. Since haploid somatic cells are generally smaller than isogenic diploids in animals (Gibeaux et al., 2018; Menon et al., 2020; Yaguchi et al., 2018), haploid embryos have a higher demand for cell proliferation (i.e., need more cells) than diploids for achieving normal organ size. Indeed, such compensatory cell number increase occurs in some cases, such as pronephric tubules and ducts in haploid newt larvae or cellular blastoderms of haploid fry (Edgar et al., 1986; Fankhauser, 1945). However, we found that haploid zebrafish larvae manifested frequent mitotic arrest and cell death across various organs, which blocked efficient cell proliferation during organogenesis. Mitotic defects in haploids also potentially led to the generation of aneuploid progenies. However, cell populations with irregular DNA content were not evident in flow cytometric analyses of haploid larvae (Fig. S1), suggesting that abnormal chromosome segregation ultimately led to cell death or growth arrest in haploids. Resolution of mitotic arrest by SAC inactivation or suppression of apoptosis by p53 depletion modestly improved organ growth in haploid larvae. These results support the idea that poor organ growth in haploid larvae stems, at least in part, from haploidy-linked defects in cell proliferation control, which abolishes the compensatory cell number increase required for ontogenesis in the haploid state.

The relatively small extent of the rescue in tissue growth by p53 depletion or SAC inactivation may be partially explained by the incomplete resolution of cell death or mitotic defects, respectively, by these treatments. However, this limited rescue also indicates that unaddressed cellular defects other than cell death or mitotic defects potentially contribute to organ growth retardation in haploid larvae, which should be clarified in future studies. We also need to note that defining “fully-rescued” haploid larval size is difficult since the relationship between ploidy and body size differs depending on biological contexts. For example, even in haplodiplontic invertebrate species such as rotifers, normally developing haploid larvae are less than half in their body length compared to their diploid counterparts (Ricci and Melone, 1998). Normally developing haploid male larvae are also smaller than diploid males in some ant species (Yamauchi et al., 2001), while they are the same in size in other ant species (Kureck et al., 2013). Therefore, it remains to be determined whether haploid vertebrate larvae reach diploid-equivalent body size when their cellular defects are sufficiently resolved.

### Developmental stage-dependent aggravation of mitotic defects

Haploid larvae suffered severe mitotic arrest with the abnormally high mitotic index after 1 dpf (Fig. 4). Corresponding to these defects, mitotic chromosome misalignment associated with spindle monopolarization became evident and gradually aggravated after 1 dpf in haploid larvae (Fig. 5). A previous study reported that normal zebrafish embryos acquire SAC function by 4 hpf (Zhang et al., 2015), suggesting that spindle monopolarization after 1 dpf activated SAC and arrest mitotic progression. Indeed, SAC inhibition by reversine sufficiently restored mitotic progression and cell proliferation patterns in haploid larvae (Fig. 6), demonstrating that monopolar spindle-driven SAC is a cause of haploidy-linked abnormality in cell proliferation. This data also shows that SAC acquisition is normal even in the haploid state in zebrafish larvae.

The correlation between centrosome loss and spindle monopolarization indicates that haploid larval cells fail to form bipolar spindle because of the haploidy-linked centrosome loss (see below for further discussions on possible future experiments to address the causality between centrosome loss and haploidy-linked cellular and larval defects). We found that the centrosome loss progressed from 1 to 3 dpf in haploid larvae, concomitantly with the gradual reduction in cell size through continuous cell divisions without full cell growth (Menon et al., 2020). Interestingly, centrosome loss occurred almost exclusively in haploid cells whose size became smaller than a certain border (Fig. 5). Because of the cell size-coupled progression of centrosome loss, severe mitotic defects and size retardation took place mainly in the organs that remained mitotically active even after 1 dpf. Based on these results, we propose that the stage-dependent propagation of centrosome loss shapes the characteristic profile of “haploid syndrome” in zebrafish larvae.

While it is currently unknown why haploid cells lose their centrosomes only below a certain cell size threshold, a possible explanation could be the depletion of centrosomal protein pool in the cytoplasm with drastic cell size reduction. Consistent with this idea, a previous study in early C. elegans embryos proposed that the total centrosomal protein amount proportional to embryonic cell size determined the extent of protein accumulation at the mitotic centrosomes (Decker et al., 2011). Alternatively, it is also possible that other primary causes, such as the lack of a second active allele producing sufficient protein pools, induced cell size reduction and centrosome loss in parallel without causality between them. It is important to note that diploid larval cells also became smaller than the size border found in haploids, though they reached the border 1 or 2 days later than haploids (Fig. 5D). However, diploid larval cells remained free from centrosome loss even when smaller than the border. Therefore, interesting future perspectives are to address the mechanism that secures centrosome number stability upon cell size changes in diploids and to specify the cause that abolishes the mechanism in haploids.

In contrast to the later organogenetic stages, haploid embryos at 0.5 dpf possessed a normal number of centrosomes. This result demonstrates that centrioles provided by the UV-irradiated sperms are functional to fully support centriole duplication during early developmental stages before 0.5 dpf. This finding excludes the possibility that centrosome loss in later haploid larvae is merely a side effect of sperm UV irradiation for gynogenesis.

### Generality of haploidy-linked cellular defects across vertebrates

In mammalian cultured cells and embryos, centrosome loss causes chromosome missegregation and chronic mitotic delay, resulting in gradual p53 accumulation that eventually blocks cell proliferation or viability (Bazzi and Anderson, 2014; Fong et al., 2016; Lambrus et al., 2016; Meitinger et al., 2016). Therefore, it is intriguing to speculate that the haploidy-linked p53 upregulation in zebrafish larvae stems from severe centrosome loss. Interestingly, haploid mammalian cultured cells also suffer centrosome loss and chronic p53 upregulation, limiting their proliferative capacity (Olbrich et al., 2017; Yaguchi et al., 2018). Our findings demonstrate a striking commonality of the haploidy-linked cellular defects between mammalian and non-mammalian species while manifesting diverse organismal defects in the haploid state.

A limitation of our current approach to alleviating mitotic arrest using reversine is that SAC inactivation does not rescue chromosome missegregation caused by premature mitotic exit with erroneous kinetochore-microtubule attachments (Santaguida et al., 2010). This precludes us from investigating the relationship between mitotic fidelity and developmental defects in haploid larvae. In a previous study, drastic mitotic defects caused by the knockdown of several key centrosomal components resulted in a similar profile of developmental defects to haploids, including the formation of microcephaly, body size reduction, and curled tails in zebrafish larvae (Novorol et al., 2013). This indicates that the severe mitotic defects observed in haploid larvae also profoundly contribute to the formation of their morphological defects. For further investigating the causality between mitotic defects and haploid developmental abnormalities, it would be ideal to have an experimental condition that restores intact centrosomal control in haploid larvae. Previously, we found that artificial re-coupling of the DNA replication cycle and centrosome duplication cycle by delaying the progression of DNA replication resolved chronic centrosome loss and mitotic defects in human haploid cultured cells (Yoshizawa et al., 2020). Though we tried to restore centrosome loss by treating aphidicolin in haploid larvae, severe toxicity of the compound on haploid larvae after 1 dpf precluded us from testing its effect on centrosome control. Genetic manipulation of centrosome duplication control in haploid larvae may provide an excellent opportunity to investigate the causality of centrosome loss in the haploid syndrome in future studies. Efforts to specify factors enabling centrosome restoration are currently underway in our laboratory.

Developmental incompetence of haploid larvae is likely a crucial evolutionary constraint of the diplontic life cycle (Sagi and Benvenisty, 2017). Though parent-specific genome imprinting would serve as the primary mechanism for blocking the development of haploid embryos in mammals, what precludes haploid development in non-mammalian species has been unknown. Based on our results, we propose that the ploidy-centrosome link, as a broadly conserved mechanism, contributes, at least in part, to limiting the developmental capacity of haploid embryos in non-mammalian vertebrates. It is an intriguing future perspective to address how the ploidy-centrosome link is preserved or modulated in invertebrate animal species, especially those with a haplodiplontic life cycle.

## Material and Methods

### Zebrafish strain and embryos

Wild-type zebrafish were obtained from the National BioResource Project Zebrafish Core Institution (NZC, Japan) or a local aquarium shop (Homac, Japan). The *Tg(fli1:h2b-mCherry)/ncv31Tg* line (Yokota et al., 2015) was provided by NZC. Transgenic animal experiments in this study were approved by the Committee on Genetic Recombination Experiment, Hokkaido University. Fish were maintained at 28.5°C under a 14 h light and 10 h dark cycle. For collecting sperm for *in vitro* fertilization, whole testes from a single male were dissected into 1 mL cold Hank’s buffer (0.137 M NaCl, 5.4 mM KCl, 0.25 mM Na_2_HPO_4_, 0.44 mM KH_2_PO_4_, 1.3 mM CaCl_2_, 1.0 mM MgSO_4_ and 4.2 mM NaHCO_3_). For sperm DNA inactivation, sperm solution was irradiated with 254 nm UV (LUV-6, AS ONE) at a distance of 30 cm for 1 min with gentle pipetting every 30 seconds (s). For insemination, we added 500 μL sperm solution to ∼200 eggs extruded from females immobilized by anesthesia (A5040, Sigma-Aldrich; 0.08% ethyl 3-aminobenzoate methanesulfonate salt, pH 7.2). After 10 s, we added 500 mL Embryo medium (5 mM NaCl, 0.17 mM KCl, 0.33 mM CaCl_2_, 0.33 mM MgSO_4_ and 10^−5^% Methylene Blue). After the chorion inflation, embryos were grown in Embryo medium at 28.5°C until use.

To inhibit pigmentation, we treated embryos with 0.03 g/L N-phenylthiourea (P7629, Sigma Aldrich) at 0.5 dpf. Dechorionation and deyolking were done manually in the cold fish ringer’s buffer without Ca^2+^ (55 mM NaCl, 1.8 mM KCl, 12.5 mM NaHCO_3_, pH 7.2). In the case of inducing p53 upregulation for checking the specificity of the anti-p53 antibody, diploid larvae were irradiated with 254 nm UV at a distance of 10 cm for 3 min at 66 hours post-fertilization (hpf). For SAC inactivation, larvae were treated with 5 μM reversine (10004412, Cayman Chemical) from the time points described elsewhere. Treatment with 0.5% DMSO was used as vehicle control for reversine treatment.

### Antibodies

Antibodies were purchased from suppliers and used at the following dilutions: mouse monoclonal anti-α-tubulin (1:800 for Immunofluorescence staining (IF); YOL1/34; EMD Millipore); mouse monoclonal anti-β-tubulin (1:1000 for Immunoblotting (IB); 10G10; Wako); mouse monoclonal anti-centrin (1:400 for IF; 20H5; Millipore); rabbit monoclonal anti-active caspase-3 (1:500 for IF; C92-605; BD Pharmingen); rabbit polyclonal anti-p53 (1:1000 for IB; GTX128135; Gene Tex); Alexa Fluor 488-conjugated rabbit monoclonal anti-phospho-histone H3 (pH3) (1:200 for IF and 1:50 for flow cytometry; D2C8; Cell Signaling Technology); and fluorescence (Alexa Fluor 488 or Alexa Fluor 594) or horseradish peroxidase-conjugated secondaries (1:100 for IF and 1:1000 for IB; Abcam or Jackson ImmunoResearch Laboratories). Hoechst 33342 was purchased from Dojinjo (1 mg/mL solution; H342) and used at 1:100.

### Flow cytometry

For isolating whole-larval cells, ∼ 10 deyolked embryos were suspended in a cold trypsin mixture (27250-018, Gibco; 0.25% trypsin in 0.14 M NaCl, 5 mM KCl, 5 mM glucose, 7 mM NaHCO_3_, 0.7 mM EDTA buffer, pH 7.2) for ∼15 min on ice with continuous pipetting. For 2 dpf or older larvae, 8 mg/mL collagenase P (Roche) was added to the trypsin mixture for thorough digestion. The isolated cells were collected by centrifugation at 1,300 rpm for 15 min at 4°C, fixed with 8% PFA in Dulbecco’s phosphate-buffered saline (DPBS, Wako) for 5 min at 25°C, permeabilized by adding an equal amount of 0.5% Triton X-100 in DPBS supplemented with 100 mM glycine (DPBS-G), and collected by centrifugation as above for removing the fixative. For DNA content and mitotic index analyses, cells were stained with Hoechst 33342 and Alexa Fluor 488-conjugated anti-pH3, respectively, for 30 min at 25°C, washed once with DPBS, and analyzed using a JSAN desktop cell sorter (Bay bioscience).

### Immunofluorescence staining

For staining active caspase-3, or pH3, larvae were fixed with 4% PFA in DPBS for at least 2 h at 25°C, followed by partial digestion with cold trypsin mixture for 3 min. For staining centrin and α-tubulin, larvae were fixed with 100% methanol for 10 min at −20°C. Fixed larvae were manually deyolked in 0.1% Triton X-100 in DPBS and permeabilized with 0.5 or 1% Triton X-100 in DPBS overnight at 4°C, followed by treatment with BSA blocking buffer (150 mM NaCl, 10 mM Tris-HCl, pH 7.5, 5% BSA, and 0.1% Tween 20) for >30 min at 4°C. Larvae were subsequently incubated with primary antibodies for >24 h at 4°C and with secondary antibodies overnight at 4°C. Following each antibody incubation, larvae were washed three times with 0.1% Triton X-100 in DPBS. Stained larvae were mounted in Fluoromount (K024, Diagnostic BioSystems) or RapiClear 1.52 (RC152001, SunJin Lab).

The density of cleaved caspase-3- or pH3-positive cells in the right midbrain area was quantified by counting these signal-positive cells using the cell counter plugin of ImageJ and dividing the positive cell number by the volume of the right midbrain area (covering 2-8 confocal layers from the top of the brain as detailed in corresponding figure legends) segmented using the polygon selection tool of Image J (NIH). To estimate mitotic cell size in 1-3 dpf larvae, we measured the area of the largest confocal sections of subepidermal mitotic cells (cells located below the top two epidermal cell layers) using median 3D filters and the polygon selection tool of ImageJ. The cell contour was judged based on microtubule staining.

### Microscopy

Immunostainings of active caspase-3 or pH3 were observed on an A1Rsi microscope equipped with a 60× 1.4 NA Apochromatic oil immersion objective lens, an LU-N4S 405/488/561/640 laser unit, and an A1-DUG detector unit with a large-image acquisition tool of NIS-Elements (Nikon). For live imaging of histone H2B-mCherry-expressing larvae, the larvae were embedded in agarose gel (5805A, Takara) in E3 buffer supplemented with anesthesia and N-phenylthiourea and observed using a TE2000 microscope (Nikon) equipped with a Thermo Plate (TP-CHSQ-C, Tokai Hit; set at 30°C), a 60× 1.4 NA Plan-Apochromatic oil immersion objective lens (Nikon), a CSU-X1 confocal unit (Yokogawa), and iXon3 electron multiplier-charge-coupled device camera (Andor). Immunostaining of centrin and α-tubulin was observed on a C2si microscope equipped with a 100× 1.49 NA Plan-Apochromatic oil immersion objective lens, an LU-N4 405/488/561/640 laser unit, and a C2-DU3 detector unit (Nikon).

### Immunoblotting

Embryos were deyolked in cold-DPBS supplemented with cOmplete proteinase inhibitor cocktail (Roche, used at 2× concentration), extracted with RIPA buffer (50 mM Tris, 150 mM NaCl, 1% NP-40, 0.5% Sodium Deoxycholate and 0.1% SDS) supplemented with 2× cOmplete and centrifuged at 15,000 rpm for 15 min at 4°C to obtain the supernatant. Proteins separated by SDS-PAGE were transferred to Immun-Blot PVDF membrane (Bio-Rad). Membranes were blocked with 0.3% skim milk in TTBS (50 mM Tris, 138 mM NaCl, 2.7 mM KCl, and 0.1% Tween 20) and incubated with primary antibodies overnight at 4°C or for 1 h at 25°C and with secondary antibodies overnight at 4°C or 30 min at 25°C. Each step was followed by 3 washes with TTBS. For signal detection, the ezWestLumi plus ECL Substrate (ATTO) and a LuminoGraph II chemiluminescent imaging system (ATTO) were used. Signal quantification was performed using the Gels tool of ImageJ.

### Measurement of larval body size

Larvae were anesthetized, mounted in 3% methylcellulose (M0387, Sigma), and observed under a BX51 transparent light microscope (Olympus) equipped with a 10× 0.25 NA achromatic objective lens (Olympus) and a 20× 0.70 NA Plan-Apochromatic lens (Olympus). We measured the body length, brain width, and lateral eye area of larvae using the segmented line tool of ImageJ. Since haploid larvae were often three-dimensionally curled or bent, we measured lengths of body axes viewed from lateral and dorsal sides and used longer ones for statistical analyses. We realized that the severeness of the haploidy-linked morphological defects tended to differ among larvae from different female parents. Therefore, we used clutches of larvae obtained from the same female parents for comparative analyses of the experimental conditions.

### Morpholino injection

The morpholino used in this study were 5’ -GCG CCA TTG CTT TGC AAG AAT TG-3’ (p53 antisense) and 5’ -GCa CCA TcG CTT gGC AAG cAT TG-3’ (4 base-mismatch p53 antisense) (Langheinrich et al., 2002). One nL morpholino (dissolved in 0.2 M KCl with 0.05% phenol red at 3 mg/mL) was microinjected into haploid embryos at the 1 or 2-cell stage using FemtoJet and InjectMan NI2 (Eppendorf).

### Statistical analysis

Analyses for significant differences between the two groups were conducted using a two-tailed Student’s *t*-test in Excel (Microsoft). Multiple group analyses were conducted using one-way ANOVA with Tukey post-hoc test in R software (The R Foundation). Statistical significance was set at *p* < 0.05. *p*-values are indicated in figures or the corresponding figure legends.

## Acknowledgment

We are grateful to Kentaro Kobayashi and other members of the Nikon Imaging Center at Hokkaido University for imaging technical support, Yoshimitsu Sagara, Kuniharu Ijiro, Hiroshi Hinou, and Shin-Ichiro Nishimura for microinjectors and microscopes, Mithilesh Mishra for kind supports, Koya Yoshizawa for supporting statistical analyses, and the Open Facility, Global Facility Center, Creative Research Institution, Hokkaido University for the flow cytometer. This work was supported by JSPS KAKENHI (Grant Numbers JP19J12210 and JP21K20737 to K.Y., and JP19KK0181, JP19H05413, JP19H03219, JPJSBP120193801, JP21K19244, and JP24K02017 to R.U.), the India Alliance Wellcome Trust/Department of Biotechnology Intermediate Fellowship IA/I/13/2/501042 to S.N., the Princess Takamatsu Cancer Research Fund, the Kato Memorial Bioscience Foundation, the Orange Foundation, the Smoking Research Foundation, Daiichi Sankyo Foundation of Life Science, the Akiyama Life Science Foundation, the Hoansha Foundation, the Terumo Life Science Foundation, and the Nakatani Foundation to R.U. The authors declare no competing financial interests.

## Author Contributions

Conceptualization, K.Y., and R.U.; Methodology, K.Y., D.S., T.Me., T.Mi., T.K., S.N., and R.U.; Investigation, K.Y., D.S., and M.H.; Formal Analysis, K.Y., D.S., M.H., and R.U.; Resources, K.Y., D.S., T.Me., A.M., T.K., S.N., and R.U.; Writing – Original Draft, K.Y., and R.U.; Writing – Review & Editing, K.Y, T.Me., A.M., S.N., and R.U.; Funding Acquisition, K.Y., S.N., and R.U.

---

**Supplemental Figure 1.**
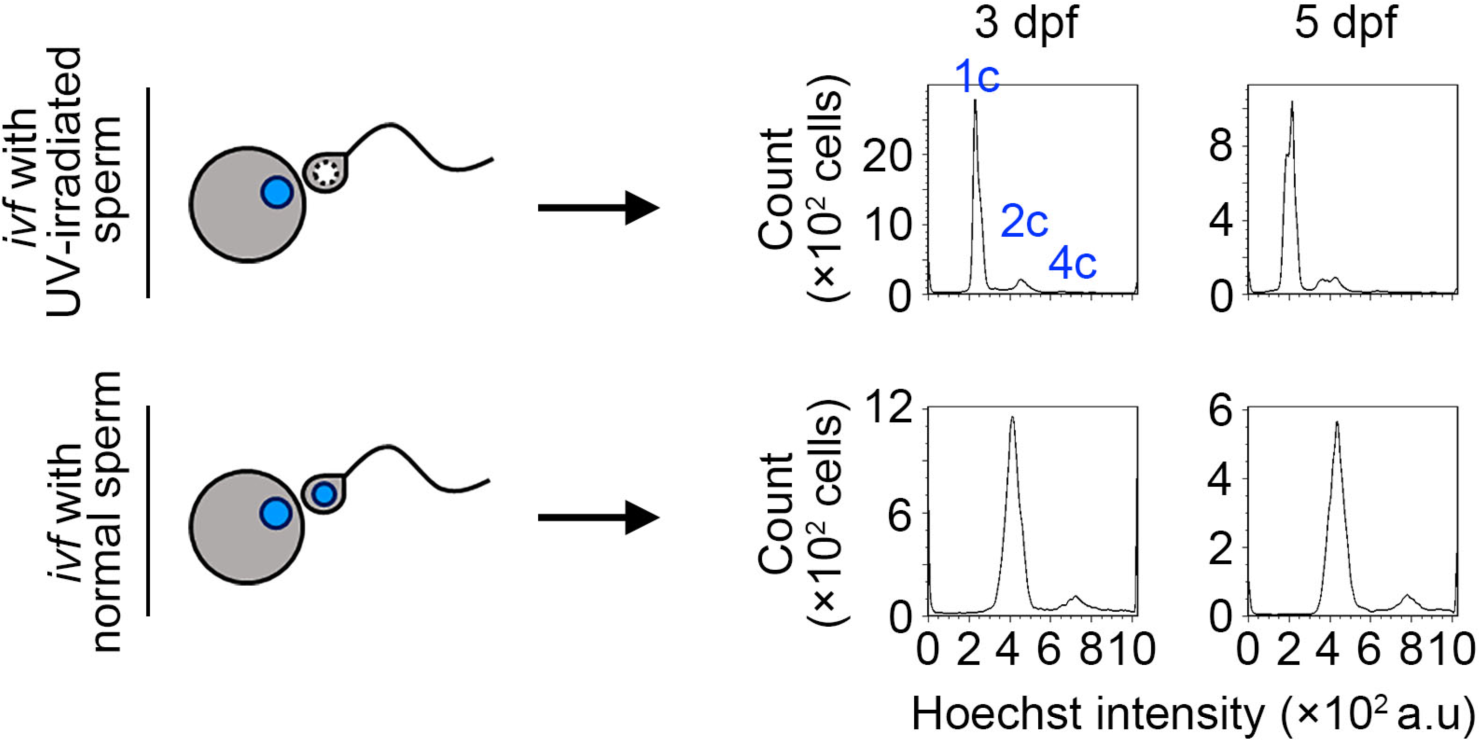
Generation of haploid and diploid larvae. Experimental scheme of in vitro fertilization for generating haploid and diploid larvae (left) and flow cytometric DNA content analysis in Hoechst-stained larval cells (right; isolated from 3- or 5-dpf larvae). The labels on the plot (1c, 2c, and 4c) indicate the relative DNA amount (c-value). Representative data from 2 independent experiments are shown.

**Supplemental Figure 2.**
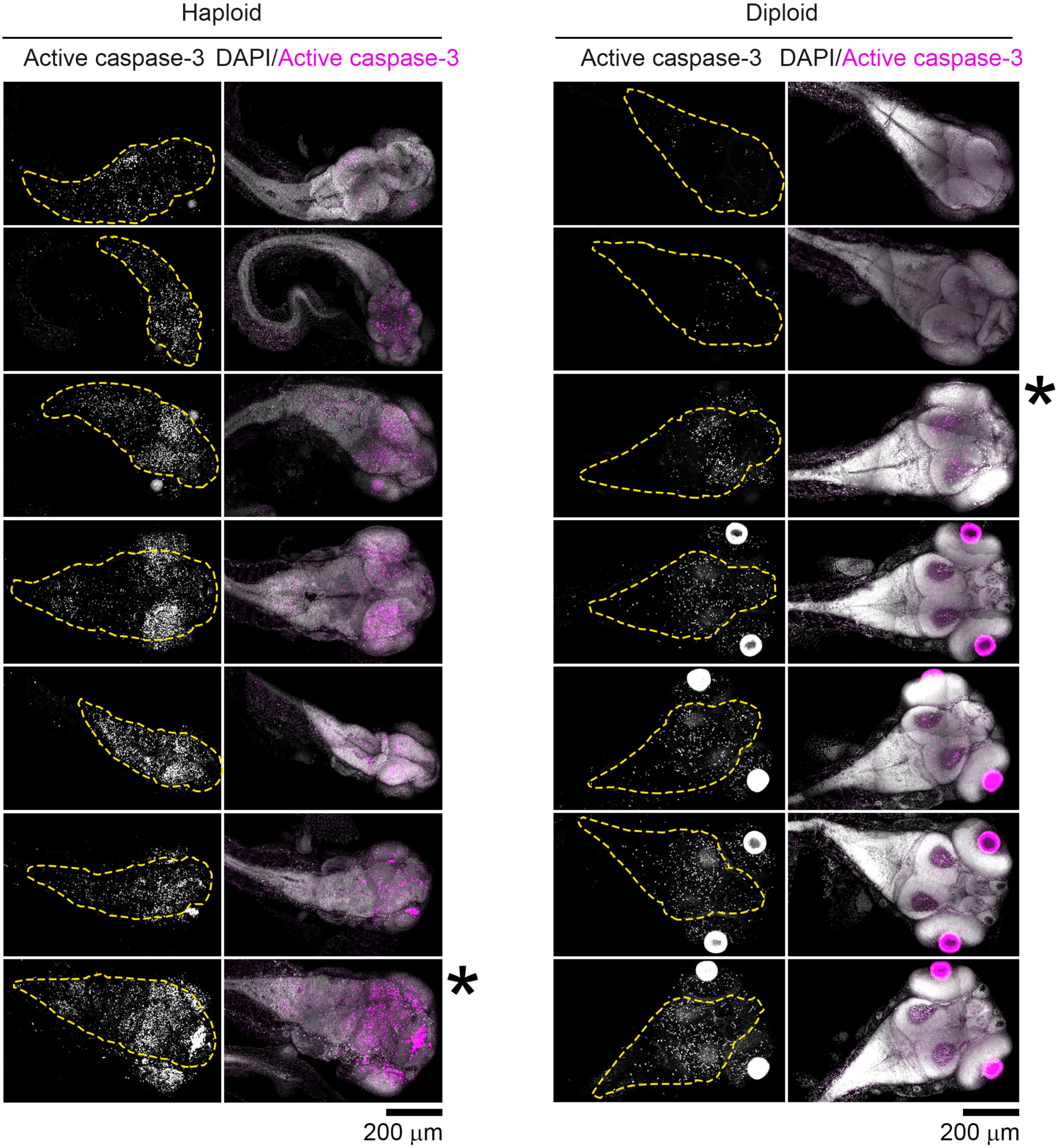
Visualization of apoptotic cells in haploid and diploid larvae. Immunostaining of active caspase-3 in whole-mount haploid and diploid larvae at 3 dpf. Z-projected images of confocal sections containing the peripheral brain surface and inner brain area. Broken lines indicate brain area. All larvae analyzed in Fig. 2C are shown. The panel includes the larvae identical to those shown in Fig. 2A (marked by asterisks).

**Supplemental Figure 3.**
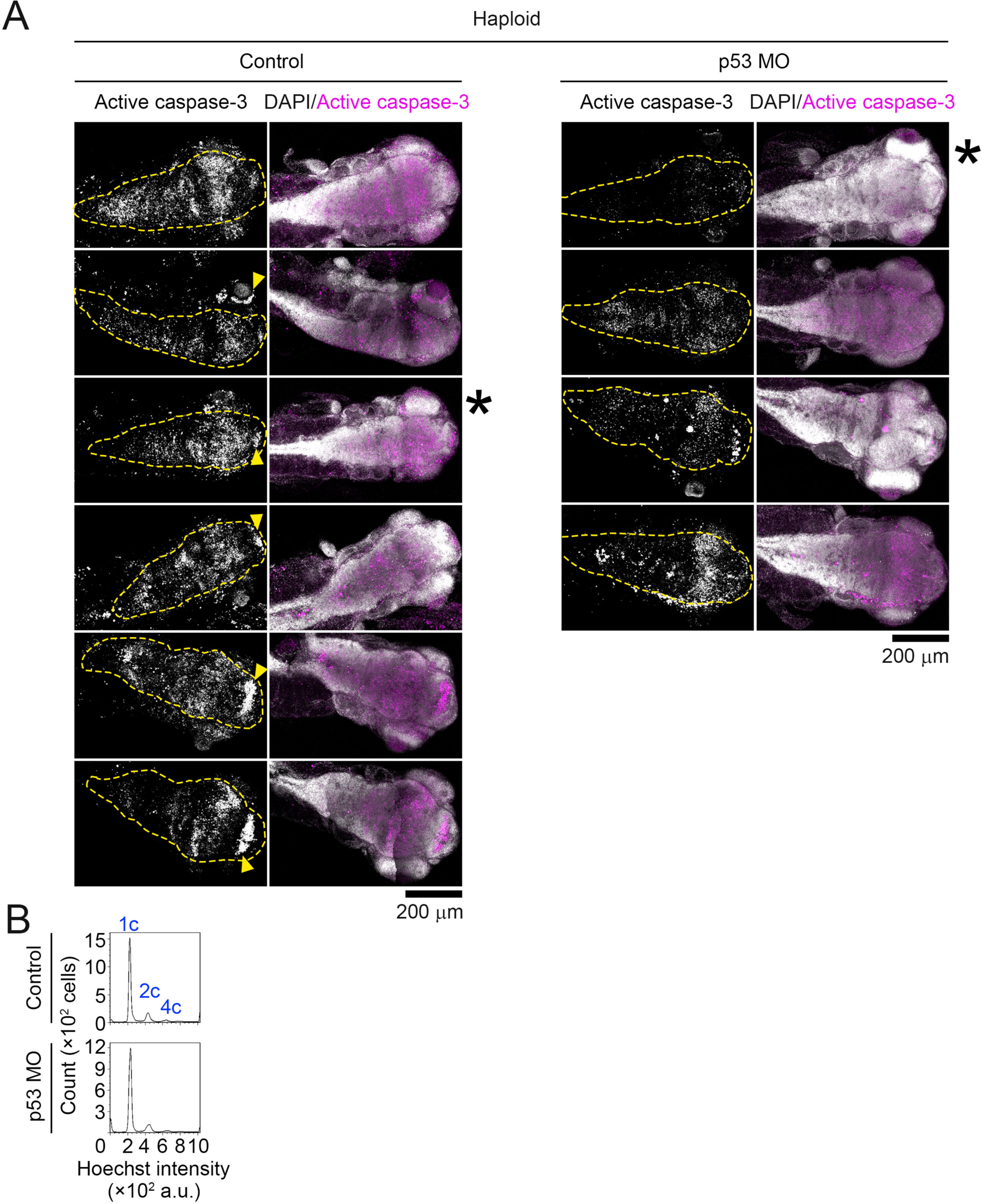
Visualization of apoptotic cells in haploid control and p53 morphants. **(A)** Immunostaining of active caspase-3 in haploid control and p53 morphants at 3 dpf. Z-projected images of confocal sections containing the peripheral brain surface and inner brain area. Broken lines indicate brain area. All larvae analyzed in Fig. 3G are shown. The panel includes the larvae identical to those shown in Fig. 3E (marked by asterisks). **(B)** Flow cytometric DNA content analysis in Hoechst-stained larval cells isolated from haploid control and p53 morphant at 3 dpf. Representative data from 2 independent experiments are shown.

**Supplemental Figure 4.**
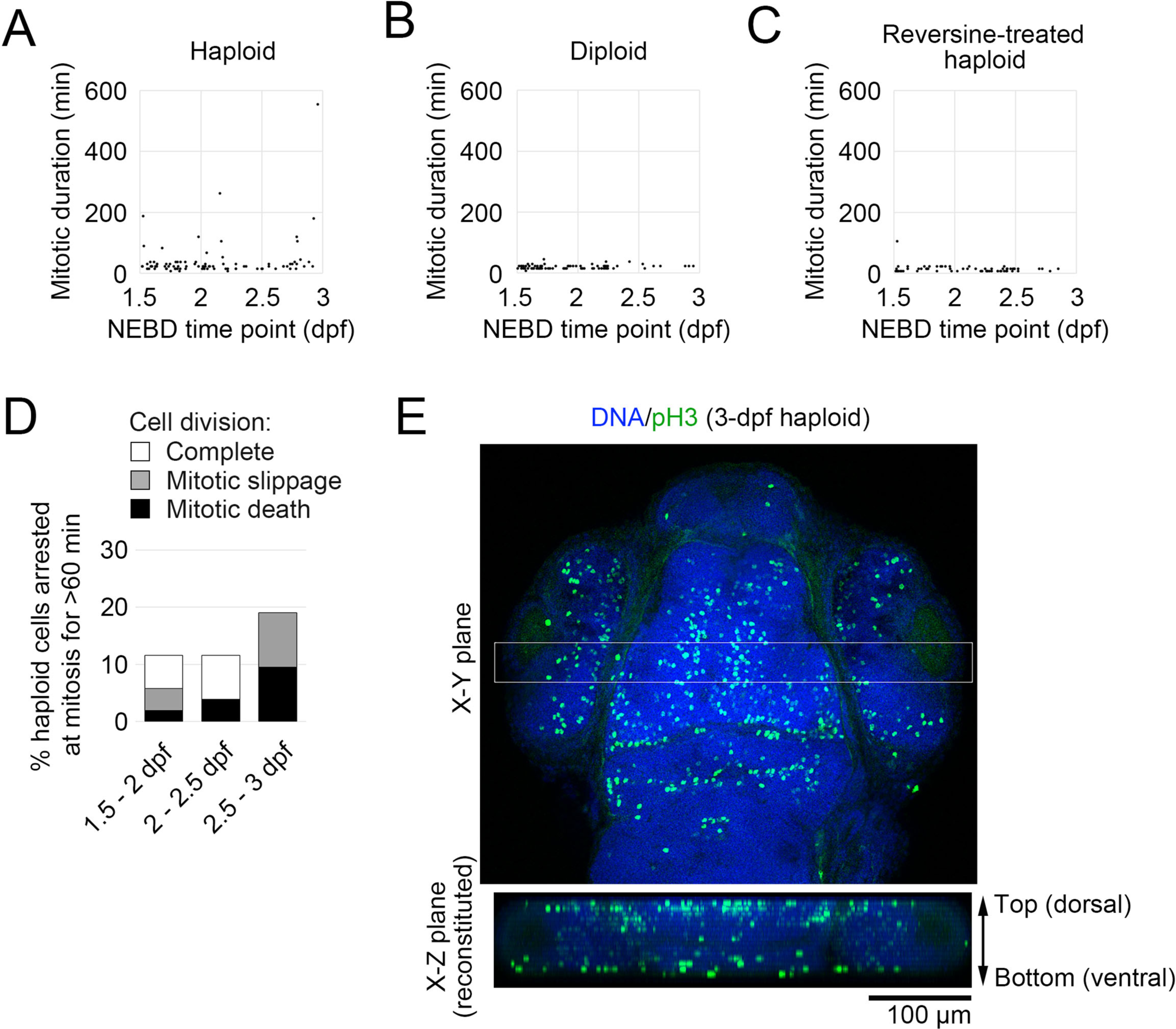
Mitotic progression in haploid, diploid, or reveresine-treated haploid larvae. **(A-C)** Mitotic length plot against NEBD time point (dpf) in haploid, diploid, or reversine-treated haploid larvae in Fig. 4C (A, B, or C, respectively). At least 74 cells of 6 larvae from 6 independent experiments were analyzed. **(D)** The frequency of mitotic fates of haploid larval cells arrested at mitosis for > 60 min (shown as percentages in the total mitotic events analyzed in A; 91 haploid larval cells of 8 larvae from 8 independent experiments). Data were sorted by NEBD time point (dpf). **(E)** A vertical section (the X-Z plane view on the bottom) reconstituted from the boxed area in the head region of pH3-immunostained 3-dpf haploid larvae. DNA was stained by DAPI.

**Supplemental Figure 5.**
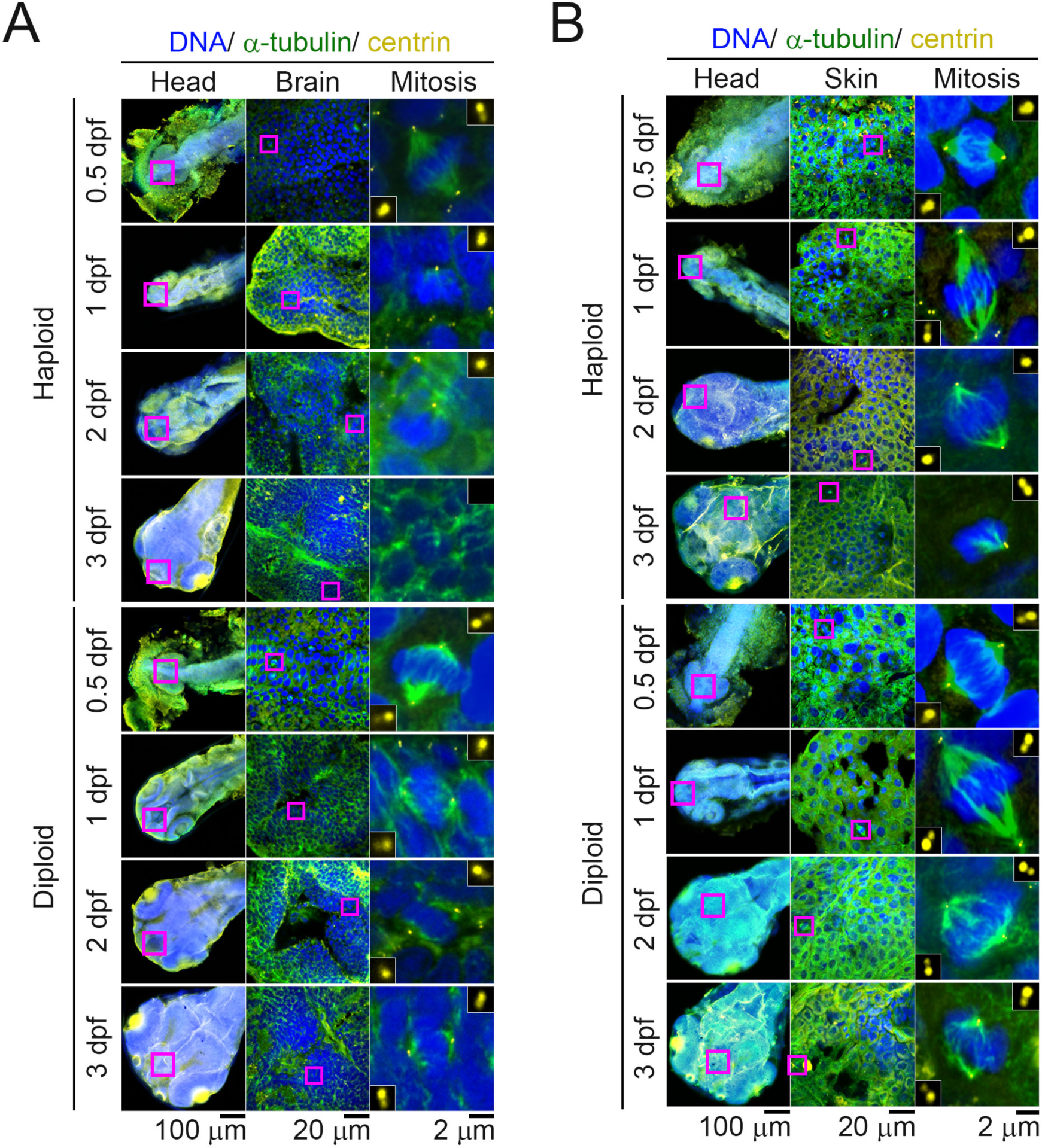
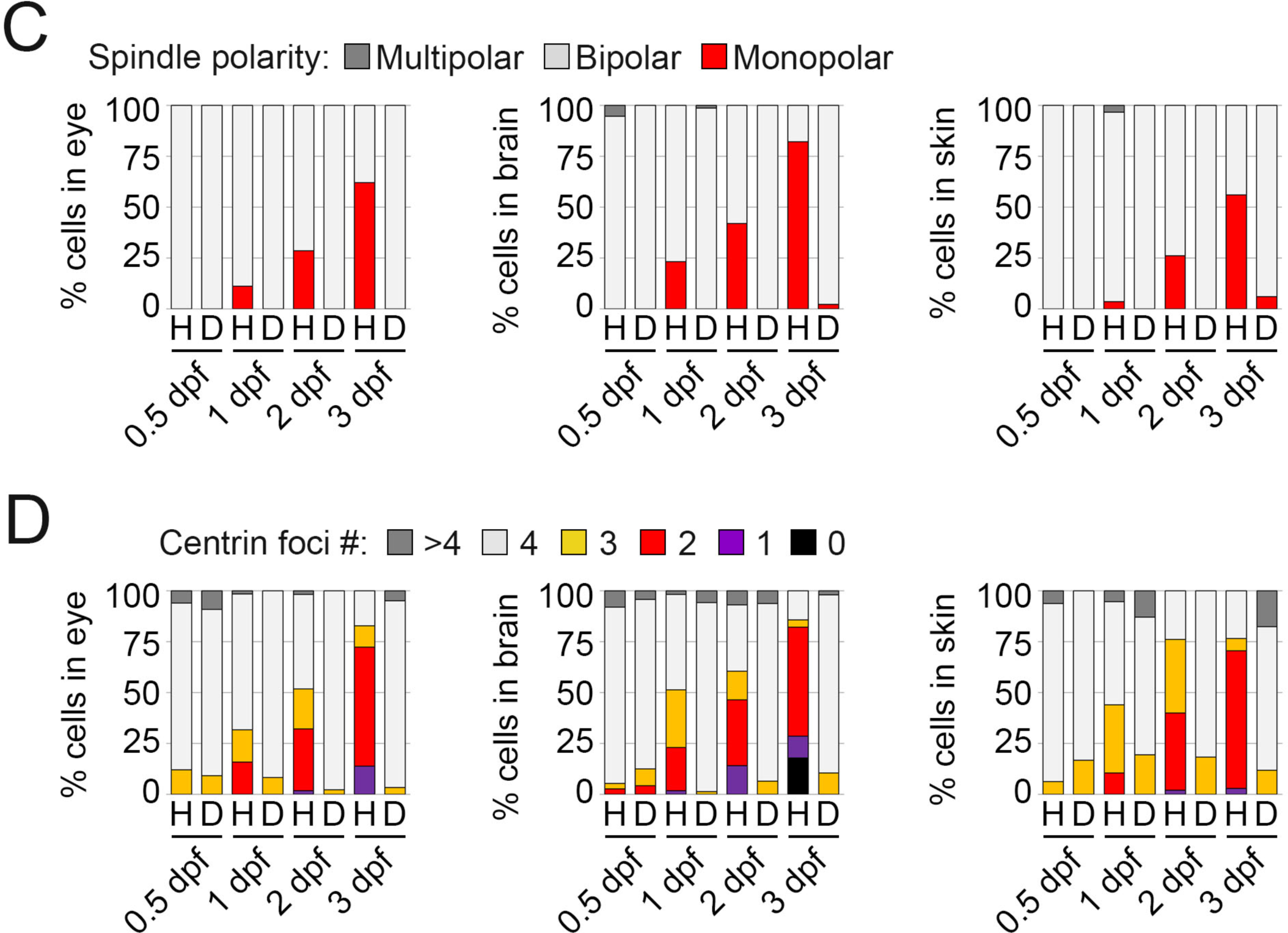
Haploidy-linked centrosome loss in different organs. **(A, B)** Immunostaining of centrin and α-tubulin in haploid and diploid larvae. Representative data of skin (A) and brain (B) of haploid and diploid larvae at 0.5, 1, 2, and 3 dpf. Magenta boxes in the left or middle panels indicate the enlarged regions of each organ (shown in middle panels) or mitotic cells (shown in right panels), respectively. Insets in the right panels show 3× enlarged images of centrioles. **(C**, **D)** Spindle polarity and centrin foci number in mitotic cells in each organ at different developmental stages. At least 11 cells of 4 larvae from 2 independent experiments were analyzed for each condition.

